# A comprehensive analysis of immune landscape of Indian triple negative breast cancer

**DOI:** 10.1101/2022.03.06.483159

**Authors:** Aruna Korlimarla, PS Hari, Jyoti Prabhu, Chanthirika Ragulan, Yatish Patil, VP Snijesh, Krisha Desai, Aju Mathews, Sandhya Appachu, Ravi B. Diwakar, BS Srinath, Maggie Cheang, Anguraj Sadanandam

## Abstract

**Background:** Triple-negative breast cancer (TNBC) is a heterogeneous disease with a significant clinical challenge. TNBC alarmingly comprises 25-30% of breast cancers in India compared with only 10-15% in the West. However, immunotherapy was approved for high-risk early-stage TNBCs in the West. Hence, a long-standing question is whether Indian TNBCs immunologically and clinically resemble Western TNBCs such that they respond similarly to immunotherapies. Here we sought to elucidate the immune landscape of Indian TNBCs for the first time, compare them to Western disease and associate them with clinical parameters, cellular types/signaling, and immunotherapy response.

**Methods:** We profiled 730 immune genes in 88 retrospective Indian TNBC samples using NanoString platform, clustered them into subtypes using a machine-learning approach, and compared them with Western TNBCs (n=422; public datasets). Subtype-specific gene signatures were identified, followed by clinicopathological, immune cell type, and pathway (multiomics) analyses. We also assessed responses to (cross-cancer) immunotherapy. Tumor-infiltrating lymphocytes (TILs) and pan-macrophage marker were evaluated using hematoxilin-eosin staining and immunohistochemistry, respectively.

**Results:** We identified three robust and similarly distributed TNBC immune transcriptome subtypes (Subtypes-1-3) in Indian women, and they are represented correspondingly in Western TNBCs and associated with well-known TNBC subtypes. Irrespective of the ethnicity, Subtype-1 harbored tumor microenvironmental and anti-tumor immune events associated with smaller tumors, younger age, and a better prognosis. Subtype-1 mainly represented basal-like/claudin-low breast cancer and immunomodulatory TNBC subtypes. Subtype-1 showed an increase in a cascade of events, including damage-associated molecular patterns, acute inflammation, Th1 responses, T-cell receptor-related and chemokine-specific signaling, antigen presentation, and viral-mimicry pathways. Subtype-1 was significantly (p<0.05) associated with pre-menopausal women, dense TILs and responses and/or improved prognosis to anti-PD-L1 and MAGEA3 immunotherapies. Subtype-2 was enriched for Th2/Th17 responses, CD4^+^ regulatory cells, basal-like/mesenchymal subtypes, and an intermediate prognosis. Subtype-3 patients expressed innate immune genes/proteins, including those representing macrophages and neutrophils, and had poor survival.

**Conclusion:** We identified three immune-specific TNBC subtypes in Indian patients with differential clinical and immune behaviors, which largely overlapped with the Western TNBC cohorts. This study suggests cancer immunotherapy in TNBC may work similarly in both populations. Hence, this may expedite pharmaceutical adoption of immunotherapy to Indian TNBC patients.

## INTRODUCTION

Breast cancer is the foremost cause of cancer-related deaths in women worldwide.^1^ Ten to fifteen percent of breast cancers in Western women are triple-negative breast cancers (TNBCs), which do not express the estrogen receptor (ER), progesterone receptor (PR), or human epidermal growth factor receptor 2 (HER2) and have a poor overall prognosis and high recurrence risk.^2^ TNBCs remain a clinical management challenge due to their aggressive characteristics and lack of targeted therapies.^3^ Chemotherapy is used to treat primary and metastatic diseases, and while targeted therapies, including immunotherapy, are slowly emerging to treat TNBCs in the West.^3^

The incidence of TNBC is 20-31% in India, which is classified as a middle-income country by The World Bank based on its per capita income, than in Western populations (high-income-countries (HIC); 10-15%).^4 5^ In India, TNBC affects younger, pre-menopausal women; and has a five-year survival rate of <30% when metastatic.^4–6^ Approximately 100,000 women are diagnosed with TNBC each year in India, with a case fatality rate of 40%, contributing to India having one of the highest breast cancer-related death rates worldwide.^4–6^ One of the proposed hypotheses for these increased death rates in India is that these TNBC patients have more aggressive disease with different immune biology than those from the West.^4 5^ However, to our knowledge, no study has explored to address this hypothesis of immune infiltration differences between Indian and Western populations. If geographically/ethnically these diseases are different, it warrants distinct treatment opportunities for TNBC in India compared to the West. Therefore, understanding the molecular heterogeneity of TNBC and the biology of its response to existing therapies in this population are essential for improving outcomes both in low-middle-income countries (LMIC) and HIC.

To characterize TNBC biology, multiple groups have defined TNBC subtypes in the West using gene expression or integrated molecular profiles.^7–9^ Immune cell infiltration into tumors is now recognized as intrinsic to tumor biology and immunotherapy responses, so a more granular analysis of immunity in TNBC is required for prediction and prognostication, especially given recent data showing that a proportion of patients respond well to immunotherapy - atezolizumab in combination with nab-paclitaxel for advanced PD-L1^+^ TNBCs (IMPassion130 study).^10^ This combination therapy has been approved in India in 2020 based on IMPassion130 study,^11^ which is no longer an approved drug in the United States (US) after the outcomes from the follow-up trial – IMPassion131 study.^12 13^ Moreover, pembrolizumab (anti-PD1 immunotherapy), combined with neoadjuvant chemotherapy, has been approved in the US for treating high-risk early-stage TNBCs (based on KEYNOTE-522 study).^14^ However, the unaddressed question is whether Indian and Western TNBCs have similar biology and clinical outcomes; therefore, combination immunotherapies from the West may benefit Indian TNBC women.

Hence, we sought to understand India-specific TNBC heterogeneity and functionally distinct immune cell types infiltrating tumors in these patients using gene expression profiling and compare this to Western TNBC patients and well-known TNBC subtypes^7–9^. The study goals were to determine: (i) whether the immune composition and their clinical associations in Indian (LMIC) TNBC women differed to that of TNBCs in Western populations (HIC); and (ii) whether differences in the cellular composition of the immune infiltrate in TNBCs were associated with pathways, immune signaling, patient survival and responses to therapy. We identified three robust immune subtypes based on consensus clustering of the expression of immune-related genes in Indian TNBC samples and found them to be represented similarly in two independent Western TNBC meta-cohorts and associated with well-known TNBC subtypes from the West. Overall, our analysis compared Indian TNBCs with those of the West at immune subtype levels (bi-directional) and suggests avenues to rationally understand the immune and clinical landscapes to potentially translate and personalize successful immunotherapy opportunities (from the West) to TNBC patients in India.

## METHODS

### Patient samples, clinical characteristics and RNA isolation

TNBC patient samples were retrospectively collected as two independent cohorts from different tertiary cancer care hospitals in Bangalore, India. Fifty out of 92 TNBC samples from one hospital, St. John’s Research Institute, were identified to be qualified for the study (**Figure 1A**). Similarly, 38 out of 48 TNBC samples were qualified for the study from Shri Shankara Cancer Hospital and Research Centre. Both the studies were approved by the respective institutional ethics review board and informed consent was obtained from the subjects.

**Figure 1.**
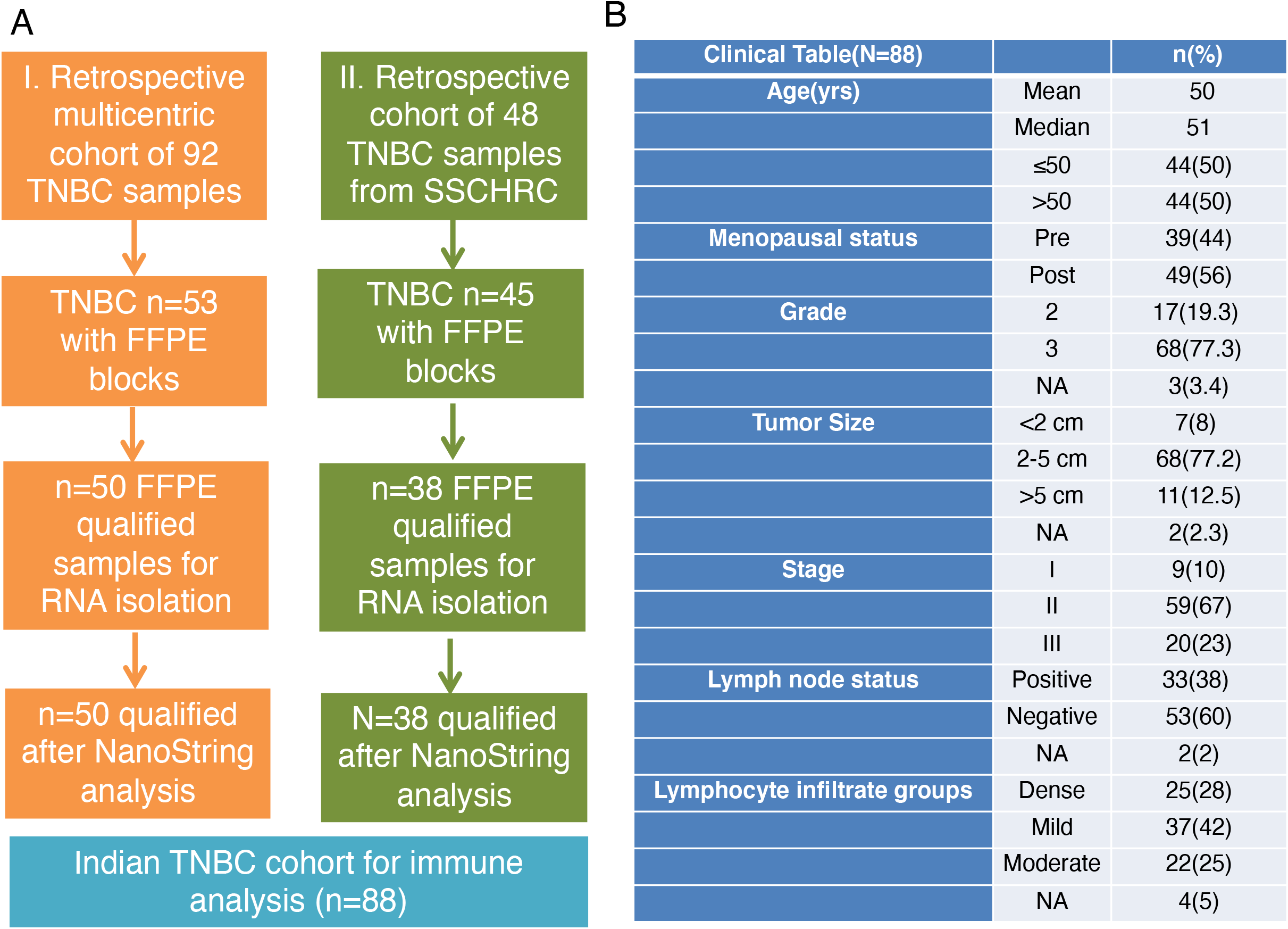
Consolidated Standards of Reporting Trials (CONSORT) diagram and clinical characteristics of Indian TNBC samples. **A**. Consort diagram showing workflow of samples selected for the study. **B**. A table of clinical characteristics of samples.

Tumor tissue samples were collected at the time of surgery (before treatment), fixed in buffered neutral formalin and processed as paraffin embedded blocks. Sections were cut from these blocks, stained with hematoxylin and eosin (H&E) and examined by a pathologist to confirm the presence of tumor. Immunohistochemistry for ER, PR and HER2 were done for determining receptor status by standard procedures^15^. Only formalin-fixed paraffin embedded (FFPE) blocks containing pre-treatment tissue with greater than 50% cancer epithelial cells were used for molecular analysis. All clinical, histopathological and demographic details were collected from the hospital medical records (**Figure 1B**). Radiologically recorded distant metastases or histologically confirmed local recurrence and death related to disease had been documented.

RNA from all FFPE specimens was extracted by the Trizol method according to the manufacturer’s instructions and as described.^16^ Briefly, 2×20 um sections were deparaffinized using xylene, washed in absolute ethanol and overnight digested with Proteinase K. Lysate was taken for extraction and RNA recovered from the aqueous phase.

### nCounter profiling and subtype analysis

nCounter Immune profiling (NanoString Technologies)^17^, subtype analysis^18^ and the PPCCA machine learning (ML) method^19^ were performed as described. A detailed description has been provided in Supplementary Methods.

### TIL Scoring

Assessment of the TILs was done according to guidelines established by the International TIL working group.^20^ Large areas of central necrosis or fibrosis are not included for evaluation, tumor was focused only on tumor-stroma at low magnification. Only mononuclear infiltrate of lymphocytes and plasma cells were included and granulocytic infiltrate in areas of tumor necrosis was not included. Based on the percentage of stromal TILs present, they were grouped into mild (no or minimal immune cells), moderate (tumor with intermediate/heterogenous infiltrate) and dense (tumor with high immune infiltrate) infiltrates.

### Immunohistochemistry and scoring

IHC was done for CD68 according to standard procedures as described in Supplementary Methods.

## RESULTS

### Clinical characteristics of the Indian TNBC cohort

To our knowledge, this is the first comprehensive immune gene, cell type, signaling and therapeutic analysis of retrospectively collected Indian TNBCs (n=88). A study flow chart and collection source are shown in **Figure 1A**. The median age of the study population was 51 years, with half ≤50 years of age and 44% pre-menopausal (**Figure 1B**). Most patients (77.3%) had grade-3 tumors, the median tumor size was 2-5 cm (77.2%), 67% had stage II disease, and 60% were lymph-node (LN) negative. Forty-two percent of samples had mild, 25% moderate, and 28% dense TILs.

### Indian TNBCs are heterogenous but can be divided into distinct subtypes with different prognoses

To characterize immune-based gene expression profiles of TNBCs in Indian women, 88 TNBC samples were profiled for 730 immune genes using the PanCancer Immune gene panel (NanoString Technologies). Batch effects in gene expression profiles between cohorts were diagnosed and corrected using our exploBATCH^19^ machine-learning (ML) tool (see Methods; **Supplementary Figure 1A-D**). After selecting 392 variable genes (standard deviation ≥1), three distinct TNBC immune gene expression subtypes were defined using unsupervised nonnegative matrix factorization (NMF)-based consensus clustering (**Figure 2A**). The robustness of the clusters and samples (or the likelihood of samples staying in the same cluster with re-iterative/consensus clustering analysis) were defined by cophenetic coefficient and silhouette statistical metrics, as previously described^18^ (**Supplementary Methods; Supplementary Figure 1E-G and Supplementary Table 1A**). The three subtypes were present in approximately equal proportions (27-38%; **Figure 2B**). 281 genes were significantly differentially expressed across the three subtypes according to supervised significance analysis of microarrays (SAM) analysis (**Supplementary Table 1B**; see Methods). Furthermore, prediction analysis for microarrays (PAM) statistical tool was used to define summary statistics for each gene per subtype (centroids; **Supplementary Table 1C; see Supplementary Methods**). Based on PAM centroids, we derived subtype-specific genes and derived scores (mean expression of genes across samples; **Figures 2C-E**). Among the 281 SAM genes, 204 were subtype-specific (non-zero) highest PAM centroid scores (**Supplementary Table 1D; Figure 2F**). A majority (79% and 19%) of 204 subtype-specific genes belonged to Subtype-1 and −2, respectively, whereas only 2% genes belonged to Subtype-3 (**Figure 2C**). Accordingly, Subtype-1/-2 showed significantly (p<0.001) higher gene diversity as measured using the Shannon Entropy method, as previously described^21^; **Figure 2G**), an index assessing immune gene expression patterns. Hence, Subtype-1 shows immune-high gene expression patterns, whereas Subtype-3 shows an immune-dormant or -exclusion pattern.

**Figure 2.**
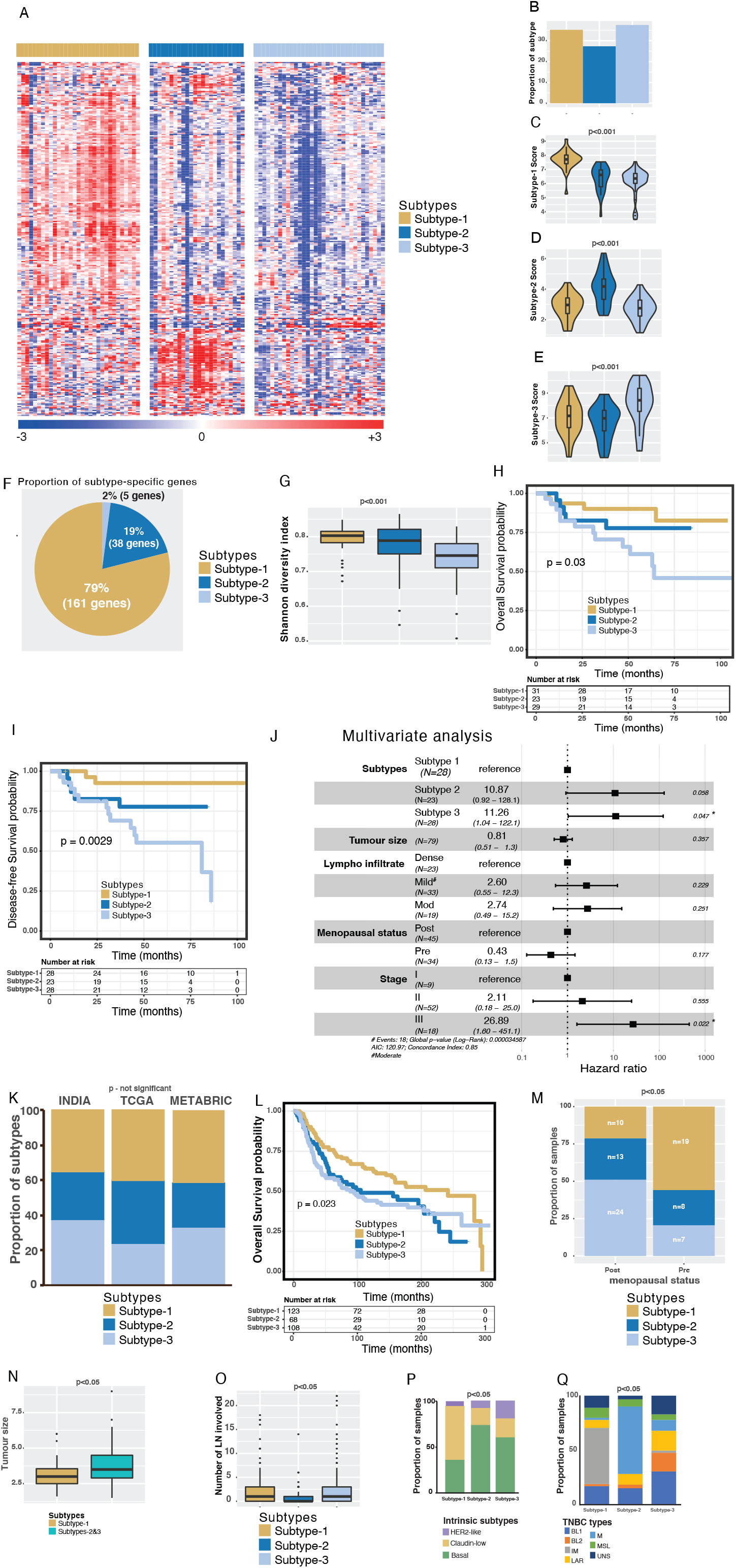
Identification and clinical characterization of immune subtypes using the Indian TNBC samples and comparison with Western TNBC cohorts. **A**. Heatmap of immune subtypes from Indian TNBC (n=88) identified by NMF clustering method. Subtypes are shown on the top bar. The scales are shown at the bottom. **B**. Proportion of immune subtypes from 88 Indian TNBCs. **C-E**. Subtype-specific scores. **F**.Proportion of subtype-specific genes out of 204 genes. **G**. Shannon diversity index of three immune subtypes. **H-I**. Kaplan-Meier curves representing H) OS (n=83) and I) DFS (n=79) from the Indian TNBC cohort. **J**. Multivariate analysis of DFS and other clinical parameters. **K**. Proportion of three immune subtypes in different cohorts of samples from India (n=88) and Western populations – TCGA (n=123) and METABRIC (n=299)**. L**. Kaplan-Meier curve representing OS from the METABRIC TNBC cohort (n=299). **M**. Proportion of pre- and post-menopausal samples represented in three immune subtypes from the Indian TNBC cohort (n=81). **N**. Tumor size differences in immune subtypes from the Indian TNBC cohort (n=86). **O**. Number of LNs involved in different immune subtypes from the METABRIC cohort (n=299). **P-Q**. Proportion of P) intrinsic and Q) Vanderbilt subtypes represented in immune subtypes from the METABRIC cohort (n=299). Kruskal-Wallis statistical test was performed for most of the analysis, except for survival analysis and proportion analysis, where the log-rank test was used for survival anlaysis and chi-squared test was used for the proportions. p-value of <0.05 was considered significant.

Subtype-3 patients had significantly (p<0.05; log-rank test) poorer overall survival (OS) and disease-free survival (DFS; **Figures 2H-I**) than Subtype-1 (favorable prognosis) and Subtype-2 (intermediate prognosis) patients. In a multivariate analysis of tumor-infiltrating-lymphocyte (TIL) groups, tumor size, menopausal status, stage, and molecular subtypes, Subtype-3 and stage-3 were independently associated with DFS (**Figure 2J**).

Next, we investigated whether these subtypes were present in TNBC samples from two Western cohorts - The Cancer Genome Atlas (TCGA, n=123)^22^ and Molecular Taxonomy of Breast Cancer International Consortium (METABRIC; n=299)^23^, which mainly represent Caucasian ethnic group from HIC. There was no significant difference in the distribution of subtypes in the Indian TNBC cohort compared with the Western cohorts, suggesting that the Indian and Western TNBCs are similar (**Figure 2K, Supplementary Table 1E-F**). There was also a similar trend in OS according to subtype in the METABRIC data as in the Indian cohort (**Figure 2L**).

Since the incidence of TNBC is reported to be higher in pre-menopausal women world-wide, including in India^2 6^, we sought to further understand the association between TNBC subtypes and menopausal status. Subtype-1 TNBCs were more common in pre-menopausal patients and Subtype-3 TNBCs were more common in post-menopausal patients (**Figure 2M**; p<0.16). A similar, but not-significant, trend of menopausal status was observed in the TCGA (p<0.16) and METABRIC (p<0.28) data (**Supplementary Figures 2A-B**).

Related to these data and as expected, Subtype-1 patients were younger (median age 44) than Subtype-2 (median age 50) and Subtype-3 (median age 54) patients (**Supplementary Figures 2C;**p<0.05, Chi-square test), and this trend (not statistically significant) was similar in TCGA and METABRIC data (**Supplementary Figures 2D-E**). Interestingly, Subtype-1 tumors were smaller than Subtype-2 and −3 tumors (**Figure 2N**), and Subtypes-1 and −3 were associated with a greater number of involved LNs (from METABRIC data) than Subtype-2 tumors (**Figure 2O**). With respect to intrinsic subtypes, Subtype-1 shows a similar distribution of claudin-low and basal-like subtypes, whereas Subtypes-2 and −3 predominantly show basal-like. Among the subtypes, Subtype-3 reveals increased HER2-like subtype, as assessed using METABRIC data (**Figure 2P**). There was no significant association between METABRIC^23^ and our subtypes (**Supplementary Figure 2F**). Interestingly, a majority of Subtype-1 is associated with Vanderbilt’s immune TNBC subtype^7^ and Subtype-2 with mesenchymal. In contrast, Subtype-3 was a mixture of all Vanderbilt’s subtypes (**Figure 2Q**). Although TCGA immune subtypes^24^ show a significant association with our immune subtypes, there is no specific association of each subtype with TCGA subtypes. Intriguingly, Subtype-2 has more than 50% samples representing wounding healing C1 TCGA immune subtypes (**Supplementary Figure 2F**). Overall, we identified three immune-specific TNBC subtypes in the Indian cohort with different immune gene expression, prognoses, and menopausal status, which largely overlapped with Western TNBC and certain molecular classifications.

### Subtype-1 is enriched for TILs, Th1 cellular immunity and a cascade of immune changes

We next investigated whether our subtypes from the Indian cohort, representing the overall TNBC immune profiles, were associated with histological assessment of TILs. In our cohort, 44% had mild, 30% had dense, and 26% had moderate TILs. Most tumors with dense TILs were Subtype-1 tumors (68%; n=25), and those with moderate (82%; n=22) and mild (81%; n=37) TIL infiltrates were mainly Subtype-2 and −3 tumors (**Figure 3A**). With respect to immune gene and cellular composition, Subtype-1 had higher expression of Th1-specific genes (*IFNG* and *IL12A*; **Figure 3B-C**), reflecting increased cellular immunity. Interestingly, most chemokines represented in the 204 subtype-specific genes were only highly expressed in Subtype-1 (**Figures 3D**). These patterns apparent in the Indian cohort were also seen in TCGA data, with increased Th1 cell, leukocyte, stromal, and TIL regional fractions and lymphocyte infiltration signature scores in Subtype-1 (**Figure 3E**). Hence, Subtype-1 appears to have a gene expression profile balanced in favor of Th1 responses and chemokine expression (**Figures 3F**).

**Figure 3.**
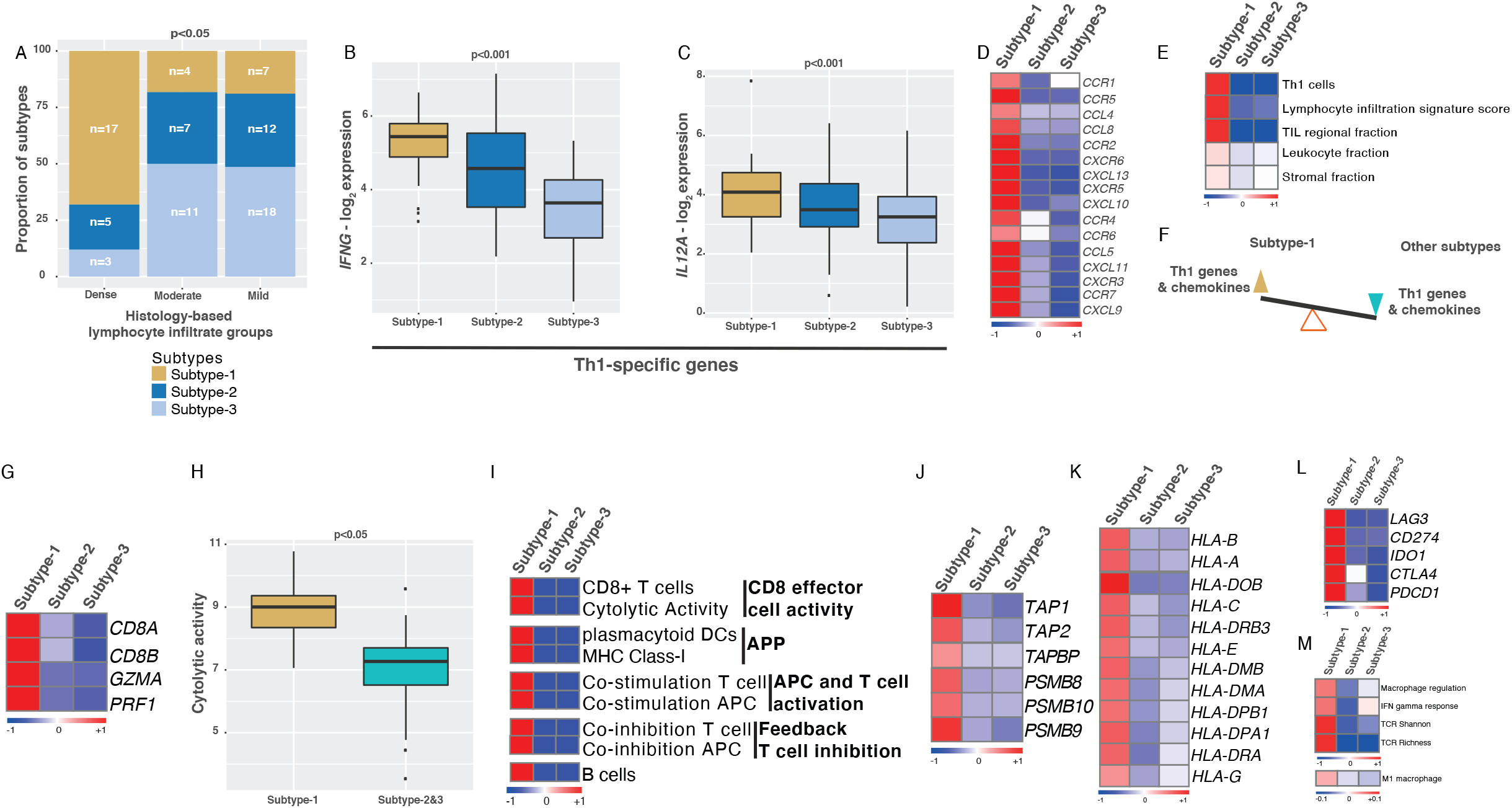
Immune and clinical characteristics specific to Subtype-1. **A**. Proportion of dense, moderate, and mild TILS represented in three immune subtypes from the Indian TNBC cohort (n=84). Chi-squared test was used for p-value calculation. **B-C**. Gene expression of Th1 response genes – B) *IFNG* and C) *IL12A* in immune subtypes from the Indian TNBC cohort (n=88). **D**. Heatmap showing chemokine expression in immune subtypes. **E**. Heatmap validation of specific immune characteristics in immune subtypes from TCGA data. **F**. Schematic representing balance in Th1 response and chemokine gene expression in subtypes. **G-L**. Heatmaps and boxplot showing changes in G) CD8 T-cell-specific genes, H) cytolytic activity, I) T and B cell types and activities based on ssGSEA analysis, J) antigen presenting and processing genes, K) MHC-I &II HLA genes, L) T-cell exhaustion genes. **M**. Heatmap showing macrophages, IFNG response, TCR diversity and intratumoral heterogeneity from TCGA data. All the figures used the Indian TNBC cohort (n=88), except figures E) and M), where TCGA data was used. Kruskal-Wallis statistical test was performed for p-value significance for those in boxplots.

We next explored the proportion and types of different immune cells within the three subtypes using single-sample geneset enrichment analysis (ssGSEA)^25^ and the entire 730 genes. The geneset distribution in the three subtypes and the inferred cell populations significantly associated (FDR <0.2) with the three subtypes are shown in **Figure 3G-K**. Subtype-1 showed increased CD8^+^ effector T-cells (and their genes *CD8A/CD8B)* and the cytolytic activity genes *GZMA and PRF1*, as assessed using cytolytic scores and ssGSEA (see Methods; **Figures 3G-I**). This increase in CD8^+^ effector T-cell activity was also associated with patterns representing increased plasmacytoid dendritic cells, major histocompatibility class (MHC) class-I, and co-stimulation of APC and T-cells and associated antigen processing and presentation (APP) genes, including genes representing transporters (TAP), immunoproteases, MHC class-II, and human leukocyte antigen (HLA) (**Figure 3I-K**). Subtype-1 also showed increased expression of immune checkpoint and T-cell exhaustion genes such as PD-1 (*PDCD1*), PD-L1 (*CD274*), *CTLA4*, and *LAG3* (**Figure 3L**), suggesting that persistent stimulation of T-cells potentially promoted T-cell exhaustion through increased expression of these genes in Subtype-1 tumors, as described^26^. Further interrogation of TCGA data showed increased INF-γ response genes and macrophage regulation, specifically anti-tumor M1 macrophages, in Subtype-1 samples (**Figure 3M**). We also examined T-cell receptor (TCR) and B cell receptor (BCR) repertoires in our subtypes using TCGA data. As expected, TCR and BCR Shannon indices and richness scores were significantly higher in Subtype-1 tumors, which might represent a highly reactive immune-associated stroma in the tumor (**Figure 3E and M**). Our subtypes capture an immune landscape and heterogeneity in TNBC beyond those represented by TIL grouping alone.

### Subtype-1 immune changes are associated with DAMPs, acute inflammation, and viral-mimicry

The cascade of adaptive immune responses and activation of dendritic cells is known to be associated with damage-associated molecular patterns (DAMPs), which are molecules released by acute inflammatory, hypoxic, or stressed cells linked to tumor progression or treatment.^27^ Hence, we examined a score derived from the average of 15 genes representing various processes involved in DAMPs.^28^ Interestingly, the DAMP gene score was significantly (p<0.05) higher in Subtype-1 tumors compared with other subtypes (**Figure 4A**). Unlike DAMPs linked to hypoxia and adaptive immunity in our previous study of pancreatic neuroendocrine tumors,^21^ DAMPs were not linked to hypoxia in Subtype-1 TNBCs, which had significantly lower hypoxia scores than Subtypes-2 and −3 as assessed using ssGSEA and the hypoxia signature from MSigDB (**Figure 4B**). Hence, we hypothesized that these significant increases in DAMP scores and Th1 responses are associated with acute inflammation^27^ in Subtype-1 tumors. As predicted, the acute inflammation gene score (mean expression of *IL15*, *IL21*, and *LTA* in each sample) was significantly (p<0.05) higher in Subtype-1 tumors compared with the other subtypes (**Figure 4C**).

**Figure 4.**
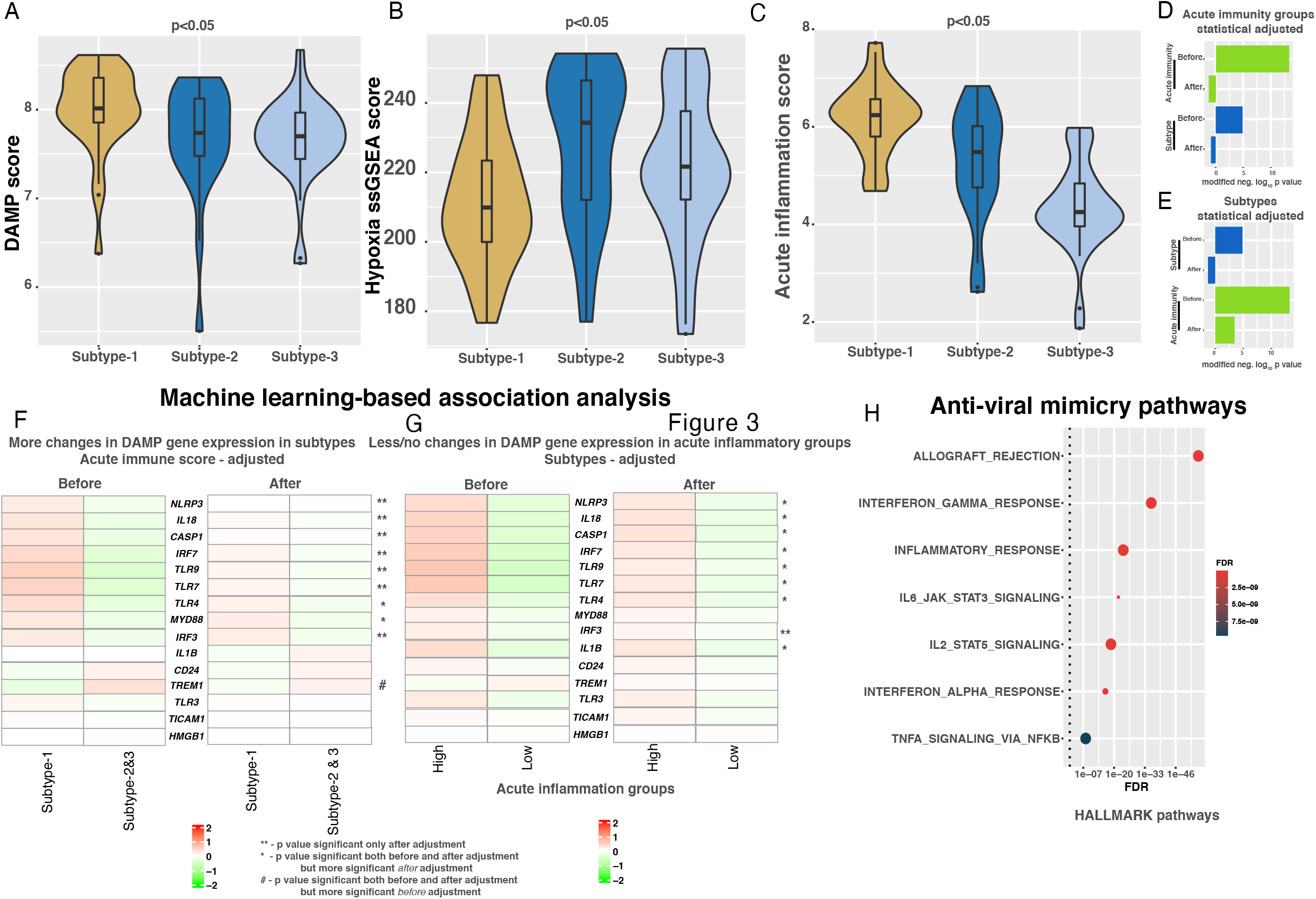
Machine-learning based association analysis of acute inflammation, subtypes and DAMP gene expression in the Indian cohort. **A-C**. Boxplots showing differential changes in DAMP, hypoxia and acute inflammation scores from the Indian TNBC cohort (n=88). Kruskal-Wallis statistical test was performed for p-value significance. **D-E**. Barplots showing significant (p<0.01; linear regression) association of 15 DAMP genes with D) acute inflammation (high vs. low) and E) subtypes (Subtype-1 vs. others) in principal component (PC1) as assessed by PPCCA method using the Indian TNBC cohort (n=88). **F**. Heatmap showing change in 15 DAMP pathway genes (median expression across samples from each subtype) in immune subtypes before and after statistically adjusting acute inflammation using the *PPCCA* method using the Indian TNBC cohort (n=88). **G**. Heatmap showing change in 15 DAMP pathway genes (median expression across samples from each subtype) in acute inflammation low and high groups before and after statistically adjusting subtypes using the *PPCCA* method using the Indian TNBC cohort (n=88). **H**. Dotplot showing enrichment of anti-viral-mimicry pathways in Subtype-1 from hallmarks gene sets from the Molecular Signature DataBase (MSigDB) database using Subtype-1-specific genes. False discovery rate (FDR) was calculated from p-values from hypergeometric test using hypeR R package (see methods).

To further explore if enrichment of DAMP genes is associated with (statistically dependent on) acute inflammation and Subtype-1, we applied our previously described ML approach, probabilistic principal component and covariate analysis (PPCCA),^19^ to the Indian TNBC cohort. This method combines probabilistic principal component and multivariate regression analyses to infer statistical associations or dependencies between three parameters (subtypes, acute inflammation, and DAMP gene expression), which cannot be assessed by Pearson or other correlation analyses. For this purpose, we categorized the severity of acute inflammation into high and low based on the median acute inflammation score as a cut-off. Then, we removed the expression differences in 15 DAMP genes between the high and low acute inflammation samples by statistically adjusting or normalizing the expression data using PPCCA. This adjusted data led to the loss of association or dependency of Subtype-1 on DAMP expression in TNBCs (p<0.05; modified negative log_10_ p-value > 0 in **Figure 4D**), as assessed using the first probabilistic principal component (pPC1; with higher proportion of variability) and linear regression analysis with the PPCCA model. Before adjustment of acute inflammation by group, both acute inflammation groups and subtypes were significantly (modified negative log_10_ p-value > 0 in **Figure 4D**) associated with DAMP expression. This was also reflected as a partial change in DAMP gene expression between Subtype-1 vs. Subtypes-2 and −3 after adjusting the acute inflammation dependency in DAMP genes (**Figure 4F**). Conversely, there was no loss of a significant association between acute inflammation and DAMP gene expression in pPC1 in the reverse analysis of adjusting subtype dependency on DAMP gene expression, followed by regression analysis of acute inflammation groups in pPC1 (**Figure 4E and G**). Moreover, we observed enrichment of host viral-infection mimicry pathways and genes in Subtype-1 (**Figure 4H**), in a similar way to our rectal cancer study^29^. These results demonstrate partial dependency of Subtype-1 on acute inflammation for DAMP enrichment and provide clues that the progression of Subtype-1 is related to viral-mimicry during infection.

### Subtype-1-specific pathways and associated immunotherapies

Pathway analysis showed enrichment of Th1, IL2, IL12, T-cell receptor (TCR), B cell receptor (BCR), cytolytic activity, PD-1, NFκB, chemokine/cytokine signaling and other related immune pathways in Subtype-1 tumors using Indian samples (**Figure 5A-B; Supplementary Table 1G-H**). These analyses allowed us to postulate a Subtype-1-specific pathway in CD4^+^ Th1 lymphocytes (based on **Figures 5A-B** and published studies^30 31^), where MEF2D pathway genes are activated through the TCR and LCK/FYN signaling (**Figure 5C**). In this pathway, CD4^+^ T-cell co-stimulation, potentially via increased chemokines/cytokines and IL2, IL12/STAT4, and LCK/FYN signaling, further triggers TNFR2-specific NFκB signaling to increase interferon-γ signaling, connecting multiple studies^32–36^. To further validate this CD4^+^ Th1 lymphocyte signaling independently, we used TCGA multi-omics profiles, including RNA-seq-based gene expression and mass spectrometry- and reverse phase protein array (RPPA)-based protein expression, and associated them with our subtypes. These independent analyses showed significant enrichment of all genes and majority of (total and/or phosphorylated) proteins from these representative pathways, suggesting activation of this pathway in Subtype-1 samples (**Figure 5D**).

**Figure 5.**
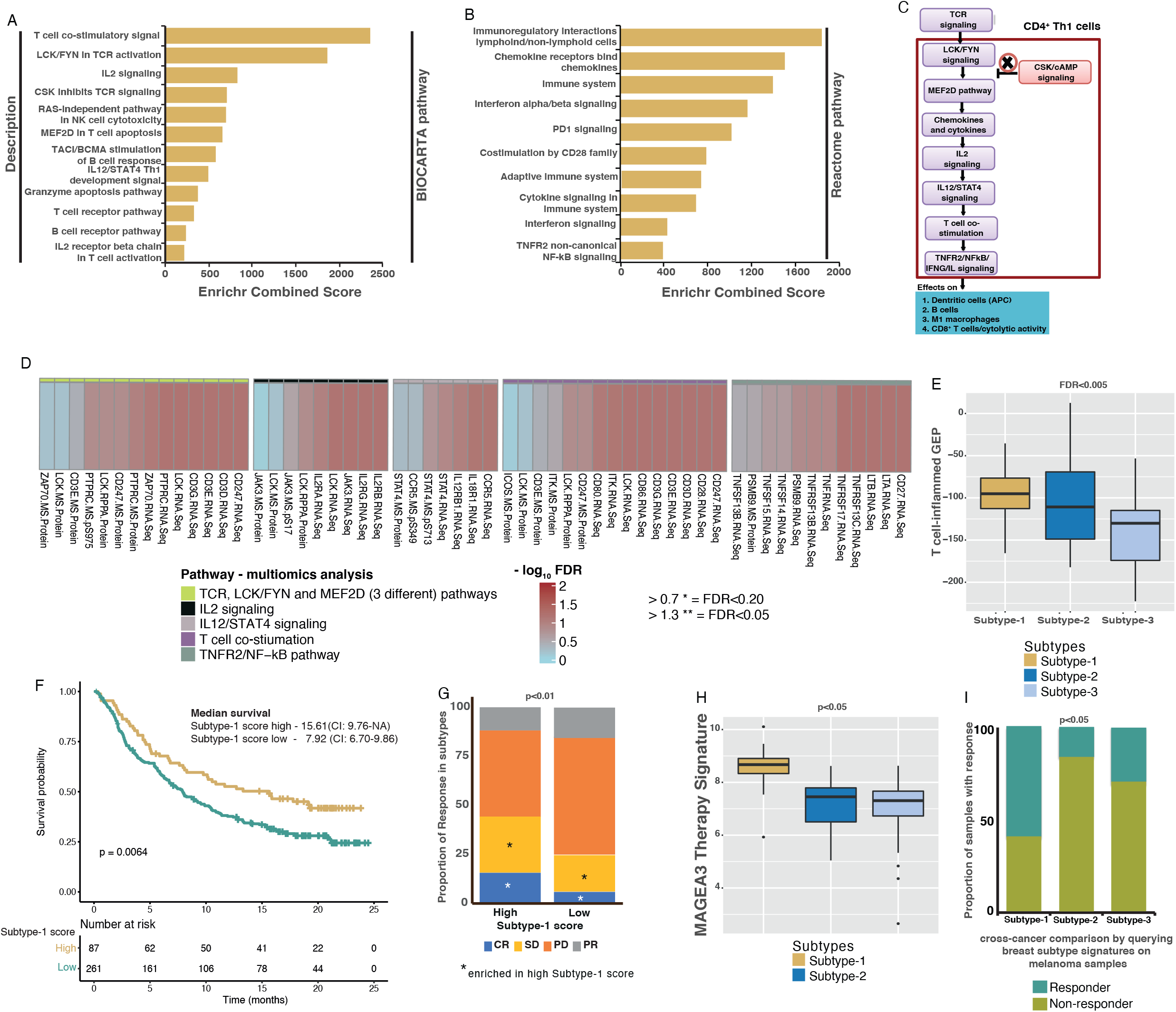
Association of gene enrichment and therapy response to immune TNBC subtypes. **A-B**. Barplot showing gene enrichment of A) BIOCARTA and B) Reactome pathways using Enrichr tool (see Methods) in Subtype-1. **C**. Schematic showing TCR and downstream signalling that effects immune cell types based on data from the Indian TNBC gene expression, pathway analysis in A-B) and literature. **D**. Multiomics enrichment analysis validation specific pathways from C) using TCGA TNBC samples (n=18). Kruskal-Wallis statistical test was performed for p-value significance. **E**. Boxplot showing differential T-cell-inflammed GEP in subtypes in the Indian cohort (n=88). **F**. Kaplan-Meier curve and median survival data showing differential OS in melanoma samples (pre-treatment; Mariathasan et al.^38^; n=348) with high and low enrichment of Subtype-1 genes and their association with immunotherapy response. Log-rank test was performed for p-value significance. **G**. Barplot showing the association of immune subtypes with clinical RECIST response to immunotherapy in melanoma (Mariathasan et al.^38^; n=298). * - represents a significant (p<0.05; Kruskal-Wallis test) enrichment in samples with high Subtype-1 score compared to those with low score. **H**. Boxplot showing differential MAGEA3 therapy response signature in subtypes in the Indian cohort (n=88). **I**. Proportion of melanoma samples (n=56) showing immune TNBC subtypes with differential MAGEA3 therapy response, as a cross-cancer comparison analysis. Kruskal-Wallis statistical test was performed for p-value significance for E) and H). Chi-squared test was performed for p-value calculations for G) and I).

Based on the above data and immune checkpoint gene expression (**Figure 3L**), we hypothesized that Subtype-1 tumors might be amenable to targeting with immune checkpoint inhibitors. Hence, we applied the expanded interferon-γ response gene signature from Ayers, et al. to our data, which has been shown to be associated with responses to anti-PD-1 therapy in multiple cancer types.^37^ We detected a significant association between the Ayers, et al., signature and Subtype-1, suggesting that Subtype-1 may respond to anti-PD-1 immunotherapy (**Figure 5E**). Moreover, we applied Subtype-1 scores to Mariathasan *et al.*, gene expression profiles, where melanoma patients were treated with anti-PD-L1 therapy.^38^ Despite the difference in cancer types, we observed that melanoma patients with high Subtype-1 scores had a better prognosis and enriched significantly (p<0.05) for compete response and stable disease than patients with low scores (**Figure 5F-G**).

Similarly, a signature derived from MAGE-A3 vaccination therapy^39^ was significantly associated with Subtype-1 tumors (**Figure 5H-I**) and, correspondingly, a majority of samples from melanoma patients responding to MAGE-A3 vaccination were significantly enriched for the Subtype-1 signature as assessed using our PAM centroids. These results suggest mechanisms of immune evasion in Subtype-1 breast tumors that may be targeted by anti-checkpoint therapy or MAGE-A3 vaccination, which warrants further assessment in TNBCs.

### Subtype-2 is associated with Th2-based humoral and Th-17 immunity

In contrast to Subtype-1, the characteristics of Subtype-2 included Th2 and Th17 gene expression, representing a shift towards humoral and Th17 immunity (**Figures 6A-E**). As expected, this Th2 response was associated with an increased proportion of CD4^+^ regulatory cells in Subtype-2 tumors (**Figure 6F**). When the tumor cell fraction was evaluated using TCGA data, Subtype-2 showed increased tumor purity (enriched for cancer cells rather than stroma) compared with the other two subtypes, opposite to Subtype-1 (**Figure 6G** and **Figure 3E**).

**Figure 6.**
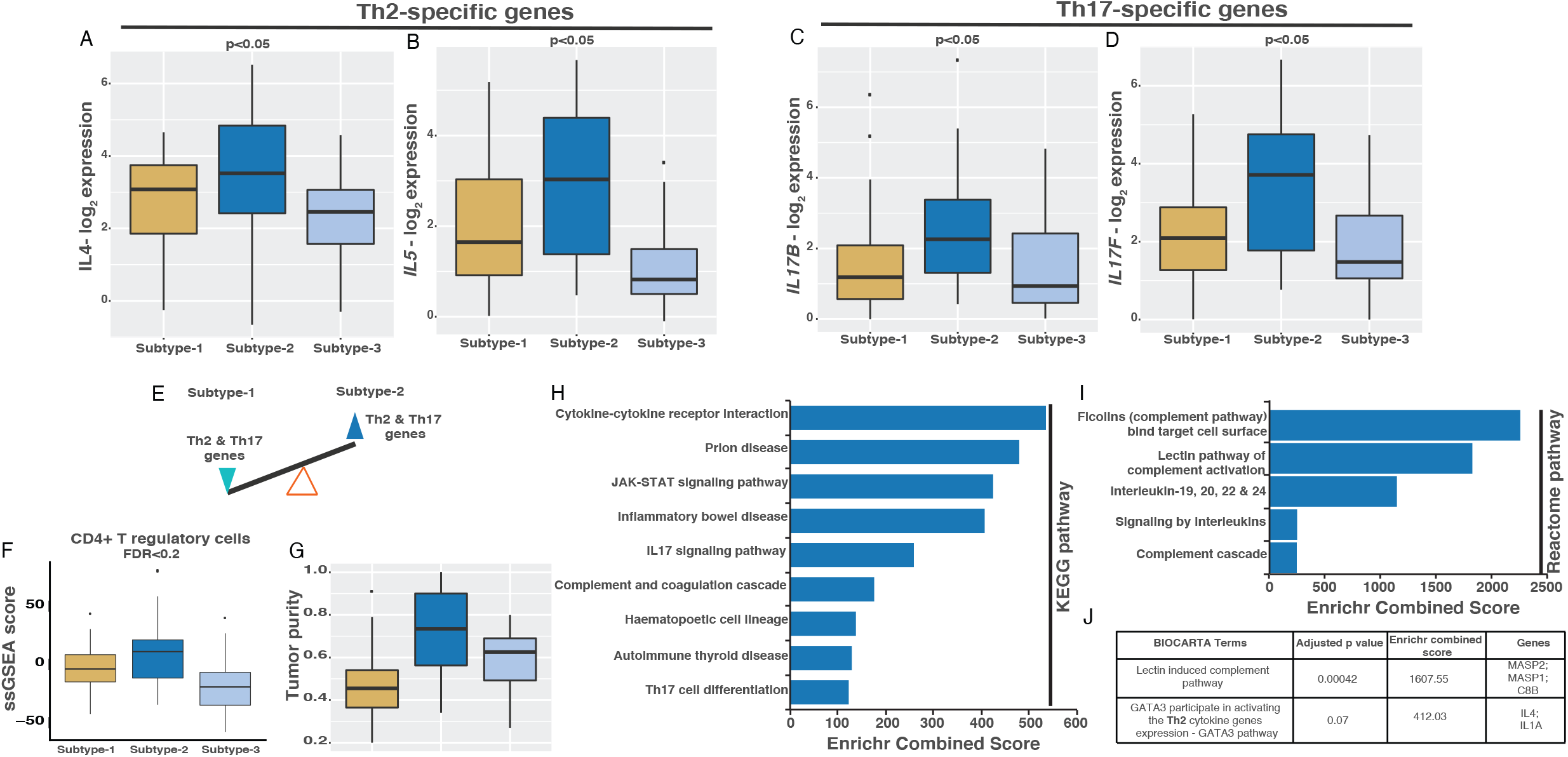
Immune characteristics specific to Subtype-2. **A-D**. Gene expression of Th2 and Th17 response genes – A) *IL4*, B) *IL5*, C) *IL17B* and D) *IL17A* in immune subtypes using the Indian TNBC cohort (n=88). **E**. Schematic representing balance in Th2 and Th17 response and chemokine gene expression in immune subtypes. **F**. Boxplot showing changes in CD4+ T regulatory cells in immune subtypes as assessed by ssGSEA analysis using the Indian TNBC cohort (n=88). **G**. Boxplot showing changes in tumor purity in immune subtypes from the TCGA data (n=101). **H-I**. Barplot showing enrichment of A) KEGG and B) Reactome pathways using Enrichr tool (see Methods) in subtype-2. **J**. A table showing BIOCARTA enrichment analysis in Subtype-2. Kruskal-Wallis statistical test was performed for p-value significance for A-G).

Next, we evaluated pathway regulation in Subtype-2 in the Indian cohort. Subtype-2 was enriched for cytokine signaling, specifically via IL17 (Th17) and lectin via the ficolin-based complement and coagulation cascade pathways (**Figure 6H-J, Supplementary Table 1I**). Also, the data suggests a balance tipping towards GATA3-induced Th2 cytokines in this subtype (**Figure 6J**). Overall, the Th2 response in Subtype-2 differs from Subtype-1.

### Subtype-3 is an immune desert subtype but enriched for innate immune cells

Subtype-3 samples significantly expressed only five subtype-specific immune genes and therefore, represents an immune desert subtype. However, there was high expression of five innate immune genes and cell types, specifically macrophage and neutrophil genes (**Figure 7A-B**). Interestingly, Subtype-3, along with Subtype-2, was relatively hypoxic compared with Subtype-1 (**Figure 4B**). We confirmed the presence of macrophages using the pan-macrophage marker CD68 in subtype-specific samples (n=39), which were indeed significantly more numerous in Subtype-3 samples (**Figure 7C-D**). CD68 positivity was confined to the tumor stroma and not epithelial nests. There was also a strong correlation between Subtype-3 and CD68-positive macrophages, with 31% of the >50% positively stained tumors belonging to Subtype-3 (p<0.05). As opposed to Subtype-1, there is an increased chronic inflammation signature in Subtype-3 patients (**Figure 7E**). The Subtype-3 signature was associated with a worse prognosis (borderline significance) in melanoma patients in Mariathasan et al. dataset^38^ (**Figure 7F**). The overall summary of all the subtypes is provided in **Figures 7G-H**.

**Figure 7.**
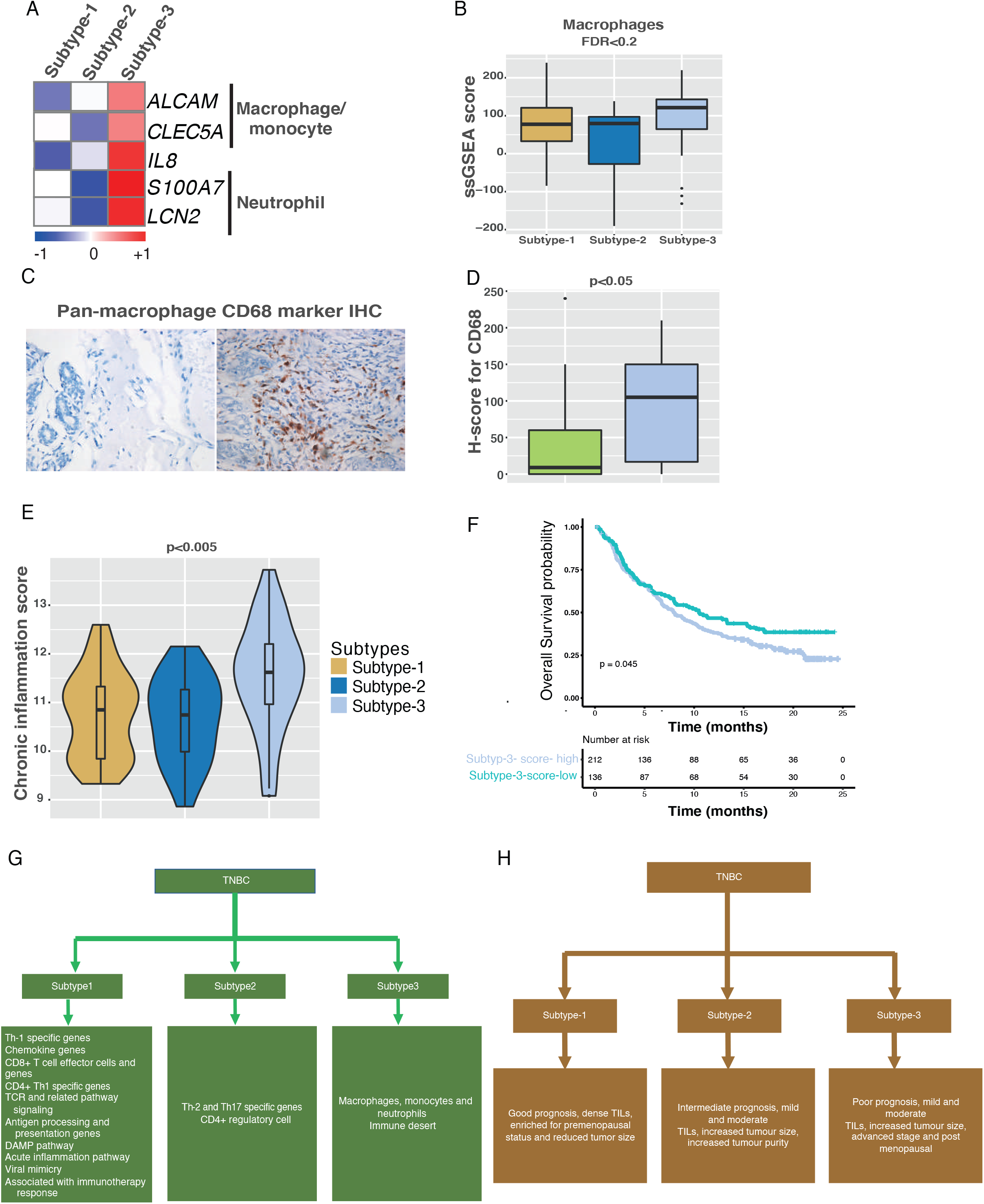
Immune characteristics and validation of macrophage markers in Subtype-3. **A**. Heatmap showing Subtype-3 specific genes associated with macrophages and neutrophils. **B**. Boxplot showing changes in macrophages in immune subtypes as assessed by ssGSEA analysis using the Indian TNBC cohort (n=88). **C-D**. IHC and quantitation (n=39) of pan-macrophage marker – CD68 in immune subtype samples using the Indian cohort. **E**. Boxplot showing differential changes in chronic inflammation scores in immune subtypes using the Indian cohort (n=88). **F**. Kaplan-Meier curve and median survival data showing differential OS in samples with high and low enrichment of immune TNBC Subtype-3 genes in melanoma samples (pre-treatment; Mariathasan et al.^38^; n=348). **G**. Molecular characteristics of subtypes. **H**. Clinical characteristics of subtypes. Kruskal-Wallis statistical test was performed for p-value significance for A), D) and E). Log-rank test was performed for p-value significance for F).

## DISCUSSION

Approximately 20-31% of all breast cancers in Indian women are TNBCs^4 5^ compared with approximately 15% in Western populations,^2^ and the incidence of TNBC is higher in pre-menopausal and young women in India.^5^ Ethnic differences in cancer prevalence and type are known to exist.^40^ For instance, about 40% of breast cancers in African American women in the US are TNBCs, a much higher prevalence than in the White ethnic groups.^41^ A better understanding of ethnicity-related disparities in cancer between Indian and Western populations would be useful for developing diagnosis and management guidelines tailored to specific populations. Given the increased prevalence of TNBC in India, there is a particular need to better understand the molecular characteristics of TNBCs in the Indian context to improve clinical outcomes by identifying potential population-specific biomarkers (if they exist) and personalizing affordable treatment regimens. Suppose the differences between Indian and Western TNBCs are minor, which is the case here, it will make it easy to adapt existing therapies, including potential immunotherapy options tailored towards TNBC, in India.

In this study, we investigated the immune landscapes of 88 breast cancer samples from South Indian women. To our knowledge, this is the first comprehensive analysis of immune transcriptome analysis of Indian TNBCs. To avoid bias, we retrospectively analyzed two distinct cohorts of retrospective samples from two different hospitals. Although these samples were collected from one major metropolitan city in South India (Bangalore), the clinical parameters were comparable to other Indian breast cancer studies, in that half of patients were aged 50 or under and over two-fifths were premenopausal.^5 6^ The age and menopausal symptom characteristics more closely resembled an African and African-American populations but were distinct from those usually seen in Western populations.^41 42^

TNBC is increasingly recognized as a heterogeneous disease with high recurrence and mortality rates that more often displays aggressive clinical features such as high grade and lymph node positivity.^2^ Despite international efforts to identify effective therapeutic targets in TNBCs by employing multi-omics strategies including proteomics, genomics, transcriptomics, and methylomics, only a handful of targeted therapies exist or under consideration for this subgroup.^3^ Breast cancer was one of the first cancer types to undergo molecular classification.^43^ Multiple groups have defined subtypes in Western TNBCs,^7–9^ and here, using Indian samples, we define three immune TNBC subtypes with distinct gene and cell type profiles and prognoses (**Figures 7G-H**). These immune subtypes defined in an Indian population were also present in Western TNBC samples and in approximately the same proportions. There is an association between our’s and Vanderbilt’s subtypes, however, our subtypes are specifically defined based on gene expression representing immune cell infiltrations that display different characteristics from Vanderbilt’s subtypes. Nevertheless, the observed geographic differences are probably less to do with the intrinsic immune biology of TNBC.

Although the incidence of TNBC is higher in young, premenopausal women,^5^ most of these women with TNBC belonged to the good prognosis Subtype-1 with active Th1 responses. In contrast, most post-menopausal women with TNBC had poor prognosis, immune desert Subtype-3 tumors. This increased risk of poor prognostic Subtype-3 in this age group may be attributable to compromised immune function, specifically dysregulated adaptive Th1 responses, with aging; the capacity to develop memory T-cells against cancer antigens is reduced in older people, compromising the ability of the immune system to reject cancer cells^44^. Therefore, subsets of pre-menopausal and post-menopausal women with TNBC may require distinct immunotherapies depending on their subtype.

Here, we have also attempted to understand the mechanistic insights of these immune subtypes associated with different immune infiltrations. The cytolytic activity of CD8^+^ T-cells seen in Subtype-1 depends on dedicated APCs, i.e., dendritic cells and, consistent with this, there was increased MHC-I expression and antigen processing and HLA genes in these tumors, which leads to co-stimulation of T-cell genes via TCR signaling in CD4 T-helper cells. Using pathway and literature^30–36^ analysis, we found that Subtype-1 was enriched for DAMP and CD4 T-helper TCR signaling, and we further validated TCR and related signaling pathways using multi-omic gene and protein data. This pathway may be associated with increased cytolytic activity in CD8^+^ T-cells, potentially reducing tumor progression and contributing to the better prognosis seen in patients with Subtype-1 TNBCs. There was also increased expression of MHC-II-associated HLA genes representing potentially activation of CD4, dendritic cells and M1 macrophages in Subtype-1 tumors, which are known to be associated with a good prognosis in several tumor types.^45^ In contrast, Subtype-3, with its low proportion of M0 and M1 macrophages and higher proportion of M2 macrophages (validated using CD68 IHC) and neutrophils, was associated with poorer outcomes. This is consistent with a study by Gentles et al.^46^, who found that macrophages, neutrophils, and plasma cells are poor prognostic markers in both breast and lung cancer.

Collectively, our findings show that there is considerable immune infiltrate variability in TNBCs, which is partly determined by the molecular characteristics of the primary tumor and that influences clinical outcomes. For example, we found that Subtype-1 may be primarily associated with an acute immune response, as described previously in other cancers^47^. Remarkably, using our PPCCA ML approach^19^, we demonstrated a significant multivariate association between acute immunity and subtypes with DAMP gene expression, suggesting increased acute immunity may be associated with or driving Th1-enriched Subtype-1 TNBCs, a hypothesis that requires validation. On the other hand, Subtype-3 may be associated with chronic inflammation and hypoxia, leading to a poorer prognosis. While the roles of Th1 responses in acute inflammation and Th2 and M2 macrophages in chronic inflammation during cancer development are well described,^47^ here we relate this to differential stratification, prognosis, signaling, and therapeutic responses. Our findings show that there is a complex relationship between acute/chronic immunity, intratumoral immune cell heterogeneity, molecular subtype, and disease progression in TNBC. Treatments that aim to enhance Th1-based cellular immunity against tumors are only effective in a subset of patients^48^, and our findings may help to identify this population using our Indian TNBC signature for specific targeting and further therapeutic development.

The success of immunotherapy in a subset of TNBC patients has raised hope of efficacy in TNBCs,^10 14^ and TNBCs with dense TILs are expected to have a better prognosis.^49^ Nevertheless, the objective response rate (ORR) for immune checkpoint inhibitors (ICI) monotherapy in metastatic TNBC is only modest, and the IMpassion130 trial for locally advanced and metastatic TNBC demonstrated that combined atezolizumab and nab-paclitaxel was more effective than monotherapy with nab-paclitaxel alone.^10^ In contrast, a follow-up trial, IMPassion131 showed no improvement in patient survival with the same therapy.^13^ Furthermore, several small studies have reported only marginal ORRs with ICIs, with poor responses likely to be multifactorial and include molecular heterogeneity, the use of ICIs as monotherapy, and their use in non-first-line settings.^3^ This is likely to be due, at least in part, to overall low PD-L1 expression (biomarker) in TNBCs,^50^ so it is essential to develop robust biomarkers to select patients who respond to immunotherapy.

Our data suggest Subtype-1 TNBC is potentially targetable with immunotherapy based on cross-cancer analysis using melanoma data. Specifically, Subtype-1 was associated with immune checkpoint inhibitor (ICI) responses and prognosis, as assessed using the anti-PD-1 treatment prediction signature from Ayers et al.^37^ and prognosis analysis using an anti-PD-L1 treated melanoma dataset.^39^ Our study also suggests an association between MAGEA3 immunotherapy and Subtype-1 tumors. Recently, a phase-III trial assessing MAGEA3 immunotherapy in stage III melanoma patients in the adjuvant setting failed.^51^ Although MAGEA3-positive cutaneous melanoma patients were selected for this trial, they were not further selected based on immune landscape such as those defined here. Similarly, a recent study has implicated the role of MAGEA3 in hepatocellular carcinoma (HCC) and suggested as a novel therapeutic avenue of targeting MAGEA3 for HCC.^52^ In contrast, It is currently uncertain what therapy would best suit patients with Subtype-2 tumors characterized by Th2/Th17 responses, infrequent LN involvement, an intermediate prognosis, and increased tumor purity; further investigation is required to identify potential immunotherapy targets associated with the immune complement cascade and GATA3. The increased wound healing response in Subtype-2 is congruent with the association of Th2 response with wound healing^53^, which might qualify them for antiangiogenic immunotherapies as described^54^. However, this subtype may benefit from bioengineered immunotherapy using collagen-binding domain (CBD)-IL12 to reactivate Th1 pathways,^55^ followed by treatment with an ICI. Subtype-3 was also associated with increased poor prognostic basal-like and HER2-like intrinsic subtypes compared with Subtype-1. Based on our data, Subtype-3 may not respond to first-line ICI-based immunotherapy.

In conclusion, here we characterized the immune heterogeneity in TNBCs in Indian women. In doing so, we identified three subtypes with distinct immune cell infiltrates, immune cell signaling, and gene signatures associated with prognosis and responses to immunotherapy. Overall, immune gene expression in Indian and Western TNBCs appears to be largely similar. This immune-transcriptome study suggests that therapies targeting immune microenvironment in Western populations may be as effective in Indian populations, however, genetic changes may need to be considered for geography-specific personalized therapy. This may accelerate pharmaceutical adoption to Indian TNBC patients.

## CONTRIBUTIONS

AS conceived and developed the idea; AS and AK further developed this idea; AK, JP and SS provided patient samples and clinical follow up data; AS, AK, HPS, CR, YP, and KD performed the experiments; AK and JP performed histopathology and related data analysis; AS, MC and SS supervised the study.

## ACKNOWLEDGEMENTS

We thank the Centre for Global Oncology at the ICR for the support.

