## Supplementary Information for "A comprehensive analysis of immune landscape of Indian triple negative breast cancer" (from another version of this preprint: version 1, 2022-03-07)

### Supplementary Methods

#### Assessment of clinical characteristics

The prognostic value of the immune subtypes was assessed using Kaplan-Meier curve, log rank test using overall survival (OS) and disease-free survival (DFS) and multivariable Cox regression using DFS.

#### nCounter Immune Profiling, batch effect and variable gene analysis

Immune gene profiling using the NanoString Technologies' nCounter® PanCancer Immune Profiling Panel and data analysis using nSolver software was carried out as per the manufacturer's instructions and as described.[1] The immune profiling for 88 Indian TNBC samples was performed in two batches based on the sources of the samples. Only those 693 genes (out of 730 genes) with zeros greater than or equal to 80% of the samples were considered. Diagnosis and correction of batch effects of the immune profiles were performed using the exploBatch machine-learning (ML) tool.[2] Highly variable 392 genes with SD>1 was selected for further analysis.

#### Other datasets and subtypes

For validation and comparison purpose, the publicly available Western cohort of only TNBC samples from TCGA[3] (123 TNBC samples; data downloaded from the Xena platform[4] in 2019) and METABRIC[5] (299 TNBC samples; data downloaded from cBio cancer genomics portal[6]) data, along with their clinical and molecular characteristics were used. In addition, multiomics data from TCGA[7] were used for subtype-specific signaling pathway analysis. For immunotherapy treatment assessment, Marthiasan et al.,[8] (n=348) gene expression and clinical dataset was used. To assess MAGEA3-based therapy, Ulloa-Montoya, et al.,[9] (n=65), gene expression data and therapy response was used. Vanderbilt et al.,[10] subtypes for METABRIC[5] were assessed using TNBCtype tool.[11]

### NMF and SAM/PAM (subtype) Analysis

NMF-based unsupervised clustering analysis was performed to identify subtypes by applying the *NMF* R package and Lee and Seung method.[12-14] The optimal number of subtypes were determined based on cophenetic co-efficient and Silhouette analysis of NMF clusters, as described[15-17]. SAM and PAM analyses were performed using the *siggenes* and *pamr* R libraries, respectively, as described.[18,19]

### Immune cell, activity, pathway and analysis

Cytolytic activity was calculated as a geometric mean of *GZMA* and *PRF1* genes as described previously[20]. Immune cell analysis was performed using ssGSEA[21] and Rooney et al.,[20] gene signatures for immune cell types. T cell-inflammed GEP was calculated as a mean expression of 18 genes from Ayer's et al.[22] KEGG[23] and REACTOME[24] pathway analyses was performed using Enrichr.[25] Acute (*IL15*, *IL21* and *LTA*) and chronic (*VEGFA*, *MIF* and *S100A8*) inflammation, DAMP (15 genes from Tang et al.[26]) and hypoxia (from Molecular Signature Database; MSigDB[27]) gene scores were calculated as mean expression per sample.

### PPCCA and multivariate gene and covariate analysis

The PPCCA method[2] was applied to assess the association between acute inflammation score, subtypes and DAMP gene expression data, as described previously by us.[28] The tool is available at https://github.com/syspremed/exploBATCH.

### Immunohistochemistry and scoring

IHC was done for CD68 according to standard procedures. Briefly, sections (5µ in thickness) were cut from FFPE blocks on poly L-lysine coated slides and subjected to deparaffinization in xylene and rehydrated in graded alcohol. After blocking endogenous peroxidase with a 3% hydrogen peroxide solution, antigen retrieval was done in 0.01M EDTA buffer at pH 8, in a heat triggered multi epitope retrieval system for immunohistochemistry from PathinSitu (Cat # MPS001). Primary blocking was done with 3% bovine serum albumin (BSA, Sigma) for 30 min at room temperature. Primary antibody for CD68(Clone KP1, Mouse monoclonal, Dako, IS609) 1 hr at room temperature. Sections were further incubated with secondary antibody (DAKO REAL™EnVision™) for 30 min as per the kit instructions, followed by development of the colour using DAB (DAKO REAL™EnVision™) for 10 min. Sections were counterstained with hematoxylin and mounted after dehydration in graded alcohol and xylene. Appropriate positive and negative controls were run for each batch. Microscopy and scoring of CD68 was done to record intensity of staining, proportion of the cells stained and localization of the protein.

### Supplementary Figure Legends

**Supplementary Figure 1. Batch diagnosis and correction and NMF analysis of 88 Indian TNBC samples. A-B.** Batch quantitative diagnosis using exploBATCH tool A) before and B) after batch correction of 88 Indian TNBC samples**. C-D.** Batch visual diagnosis using PCA C) before and D) after batch correction of 88 Indian TNBC samples. **E-G.** NMF analysis showing E) cophenetic analysis, F) Silhouette statistics and G) consensus plot of 88 Indian TNBC samples.

**Supplementary Figure 2. Menopausal, age and subtype analysis of 88 Indian TNBC samples. A-B.** Association of menopausal status with immune subtypes in A) TCGA and B) METABRIC TNBC cohorts. **C-E.** Association of age with immune subtypes in C) Indian, D) TCGA and F) METABRIC TNBC cohorts. **F-G.** Association of immune subtypes with F) integrated breast cancer subtypes and G) TCGA immune subtypes.

### Supplementary Table Legends

**Supplementary Table 1. A.** Indian TNBC samples and their immune subtype identities. **B.** SAM analysis results. **C.** PAM Centroids. **D.** Subtype specific genes. **E.** TCGA TNBC samples and their immune subtype identities. **F.** METABRIC TNBC samples and their immune subtype identities. **H.** BIOCARTA Subtype-1 pathways. **H.** Reactome Subtype-1 pathways. **I.** Reactome Subtype-2 pathways.

### Supplementary Figure 1

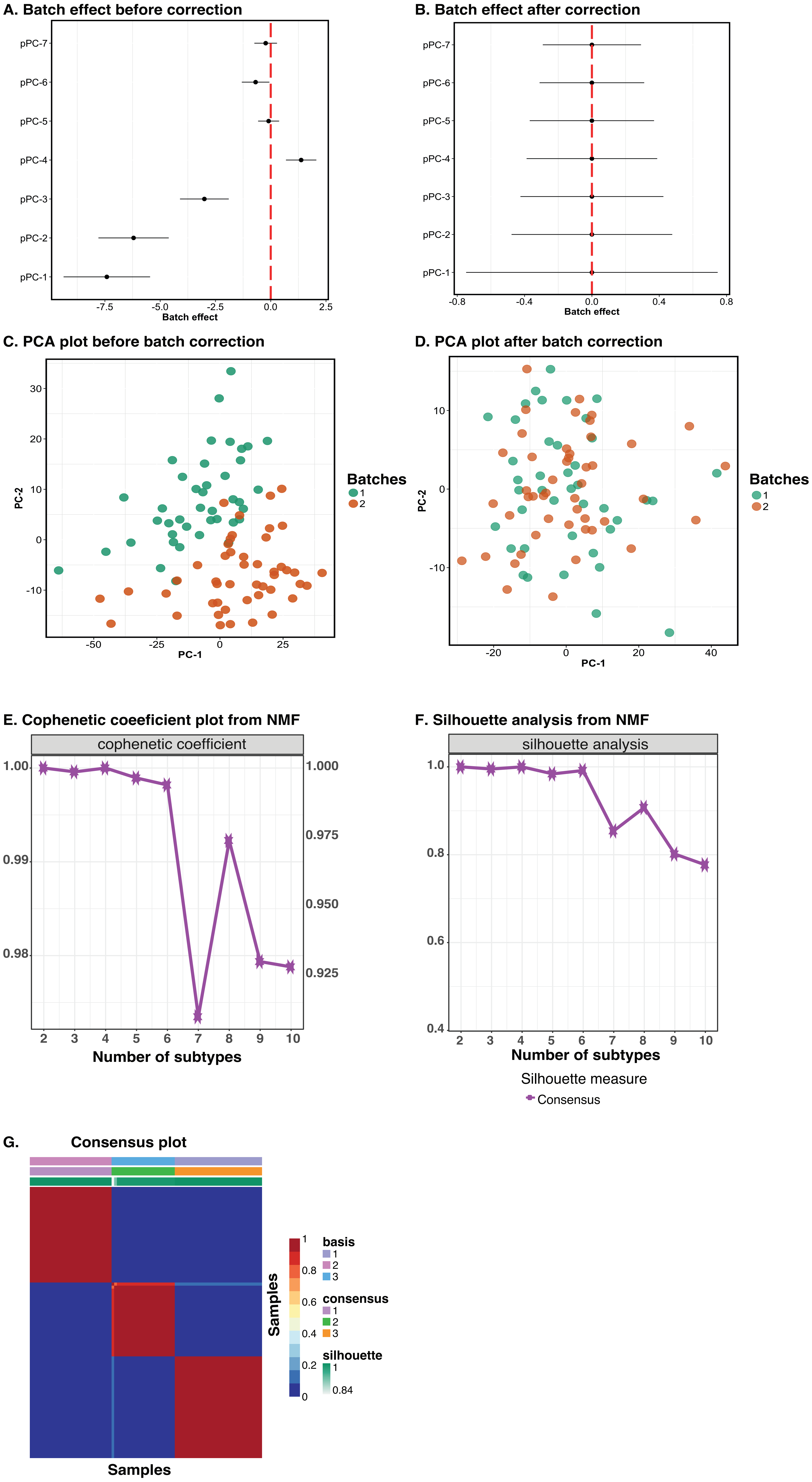

Supplementary Figure 2

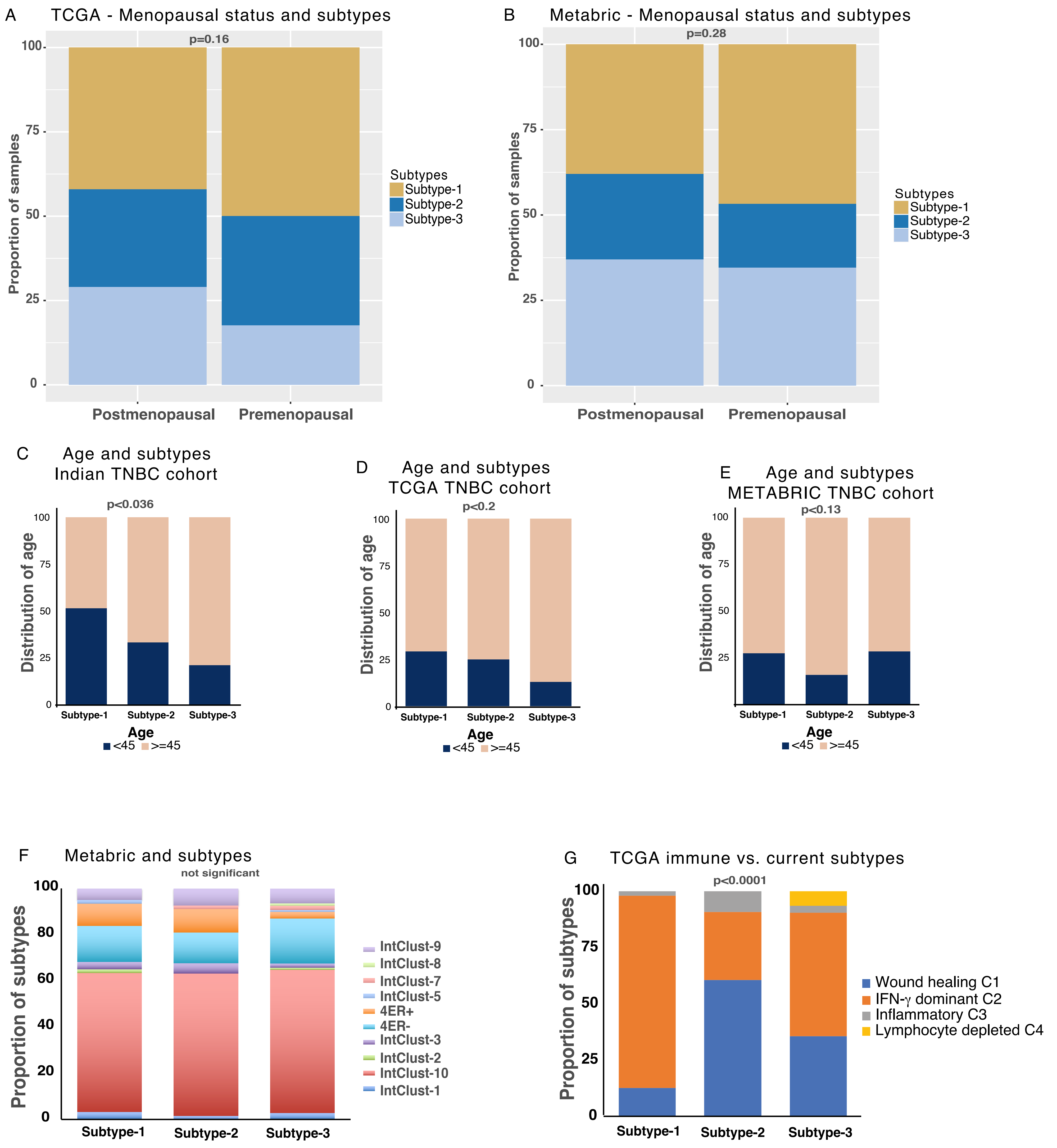
